# pysster: Classification of Biological Sequences by Learning Sequence and Structure Motifs with Convolutional Neural Networks

**DOI:** 10.1101/230086

**Authors:** Stefan Budach, Annalisa Marsico

## Abstract

**Summary:** Convolutional neural networks (CNNs) have been shown to perform exceptionally well in a variety of tasks, including biological sequence classification. Available implementations, however, are usually optimized for a particular task and difficult to reuse. To enable researchers to utilize these networks more easily we implemented pysster, a Python package for training CNNs on biological sequence data. Sequences are classified by learning sequence and structure motifs and the package offers an automated hyper-parameter optimization procedure and options to visualize learned motifs along with information about their positional and class enrichment. The package runs seamlessly on CPU and GPU and provides a simple interface to train and evaluate a network with a handful lines of code. Using an RNA A-to-I editing data set and CLIP-seq binding site sequences we demonstrate that pysster classifies sequences with higher accuracy than other methods and is able to recover known sequence and structure motifs.

**Availability:** pysster is freely available at https://github.com/budach/pysster.

## 1 Introduction

In recent years, deep convolutional neural networks (CNNs) have been shown to be an accurate method for biological sequence classification and sequence motif detection (Alipanahi *et al*., 2015) (Kelley *et al*., 2016) (Angermueller *et al*., 2017). The increasing amount of sequence data and the rise of general-purpose computing on Graphics Processing Units (GPUs) have enabled CNNs to outperform other machine learning methods, such as random forests and support vector machines, in terms of both classification performance and runtime performance (Kelley *et al*., 2016) (Angermueller *et al*., 2017). While a number of publications have made use of CNNs on biological data, these implementations are usually hard to reuse (Alipanahi *et al*., 2015) or tailored to a specific problem, such as prediction of DNA CpG methylation from single-cell data (Angermueller *et al*., 2017). Basset (Kelley *et al*., 2016) and iDeep (Pan and Shen, 2017) represent more general frameworks for training of CNNs on DNA and RNA data, respectively. However, they don’t provide detailed motif interpretations, such as motif locations and class enrichment of motifs, and they are not able to learn structure motifs in the RNA case.

To address these issues and to enable researchers to easily utilize CNNs, we implemented pysster, a python package for training CNN classifiers on biological sequences. Supervised classification is enabled by the automatic detection of sequence motifs. Our package focuses on interpretability and extends previous implementations by providing information about the positional and class enrichment of learned motifs. Moreover, by incorporating structure information, e.g. in the form of secondary structure predictions for RNA sequences, it is possible to learn structure motifs corresponding to the sequence motifs. We demonstrate that our tool is able to learn well-known motifs for an RNA A-to-I editing data set and multiple CLIP-seq data sets and that it outperforms GraphProt (Maticzka *et al*., 2014), a state of the art classifier for RNA sequences and structures, both in terms of classification and runtime performance. Providing a simple programming interface we hope that our package enables more researchers to make effective use of CNNs in classifying and interpreting large sets of biological sequences.

## 2 Implementation And Features

We implemented an established network architecture and multiple interpretation options as an easy-to-use python package. The basic architecture of the network consists of a variable number of convolutional and max-pooling layers followed by a variable number of dense layers (Figure 1A). These layers are interspersed by dropout layers after the input layer and after every max-pooling and dense layer. Using an automated grid search, the network can be tuned via a number of hyper-parameters, such as number of convolutional layers, number of kernels, length of kernels and dropout ratios. The main features of the package are:

- multi-class and single-label or multi-label classifications
- sensible default parameters and an optional hyper-parameter tuning
- learning of motifs + interpretation in terms of positional and class enrichment (Figure S1) and motif co-occurrence (Figure S2)
- support of input strings over user-defined alphabets (i.e. applicable to DNA, RNA and protein data)
- optional use of structure information, handcrafted features and recurrent layers
- visualization of all network layers using visualization by optimization (Olah *et al*., 2017)
- seamless CPU or GPU computation by building on top of TensorFlow (Abadi *et al*., 2016) and Keras (Chollet *et al*., 2015).

**Fig. 1.**
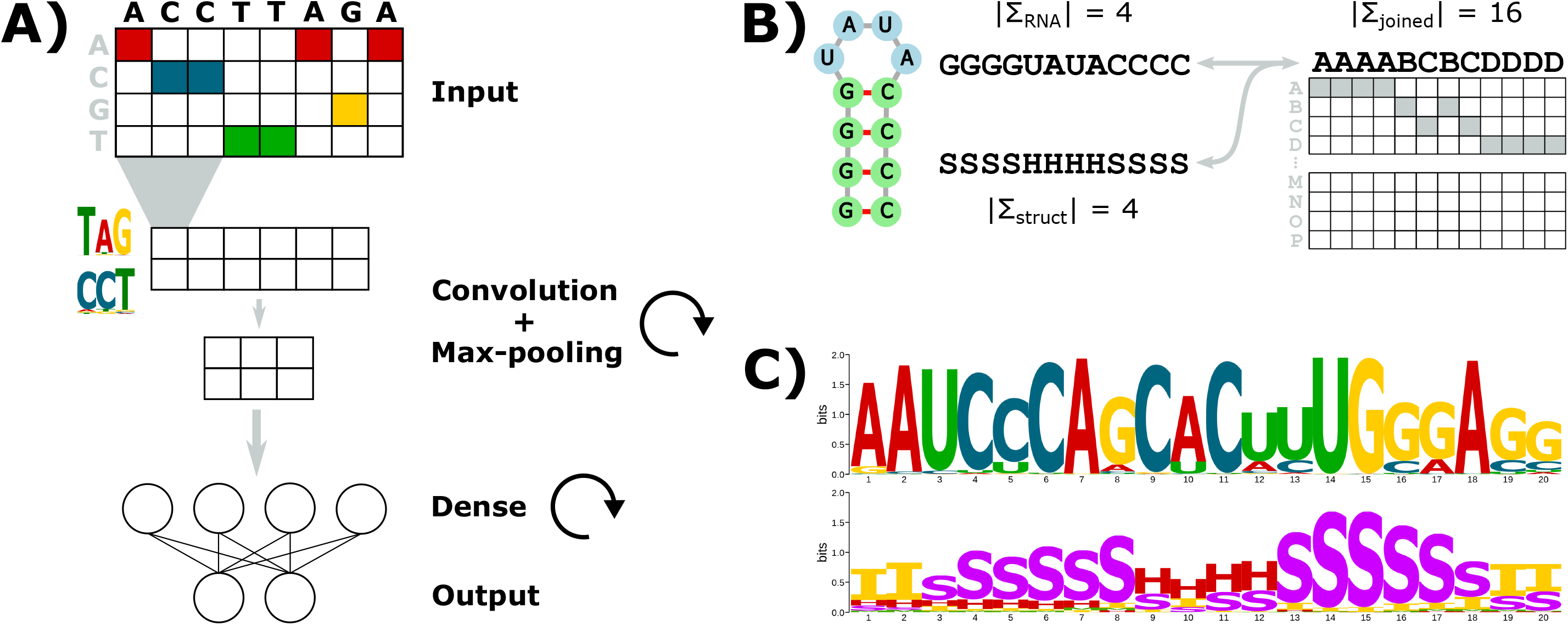
A) The basic network architecture consists of a variable number of convolutional/max-pooling stacks followed by a variable number of dense layers interspersed by dropout layers. The network can be tuned via an extensive list of hyper-parameters. The network input are one-hot encoded sequences and the network outputs predicted probabilities, indicating class membership. B) RNA sequence and structure input strings are encoded into a single string by combining the sequence alphabet and the secondary structure alphabet into an extended alphabet consisting of arbitrary characters. Subsequently, this string is one-hot encoded and used as the network input. C) For the motif interpretation the string over the arbitrary alphabet can be decoded into the two original strings to construct sequence logos for the original alphabets. The shown example motif corresponds to an ALU repeat motif found in the classification task of RNA A-to-I editing sites (see the tutorial workflow on github for detailed information).

Structure information (e.g. for RNA in the form of dot-bracket strings or annotated dot-bracket strings) is incorporated into the network by encoding the given sequence string over an alphabet of size N and the corresponding structure string over an alphabet of size M into a single new string using an extended alphabet of size N*M (Figure 1B). The network is then trained on these new strings, which can be decoded back into the original strings after the training to enable the visualization of two motifs as position-weight matrices (Figure 1C). Detailed descriptions of the model and visualization options can be found in the tutorials and documentation.

## 3 Case Studies

Pysster is freely available at https://github.com/budach/pysster. Its documentation includes a workflow tutorial that showcases the main functionality on an RNA A-to-I editing data set (Picardi *et al*., 2017). Editing sites are known to be enriched in repetitive ALU sequences and we show that we are able to classify the editing location with high accuracy and that we learn known ALU motifs (Figure 1C). Finally, we have trained our tool on sequences derived from CLIP-seq data for a number of RNA binding proteins with known binding site motifs (Table S1). Similar to GraphProt and ssHMM (Heller *et al*., 2017), an unsupervised hidden Markov model-based approach for learning sequence/structure motifs, pysster can recover the known motifs, but outperforms GraphProt both in terms of classification and runtime performance.

---

**Figure S1.**
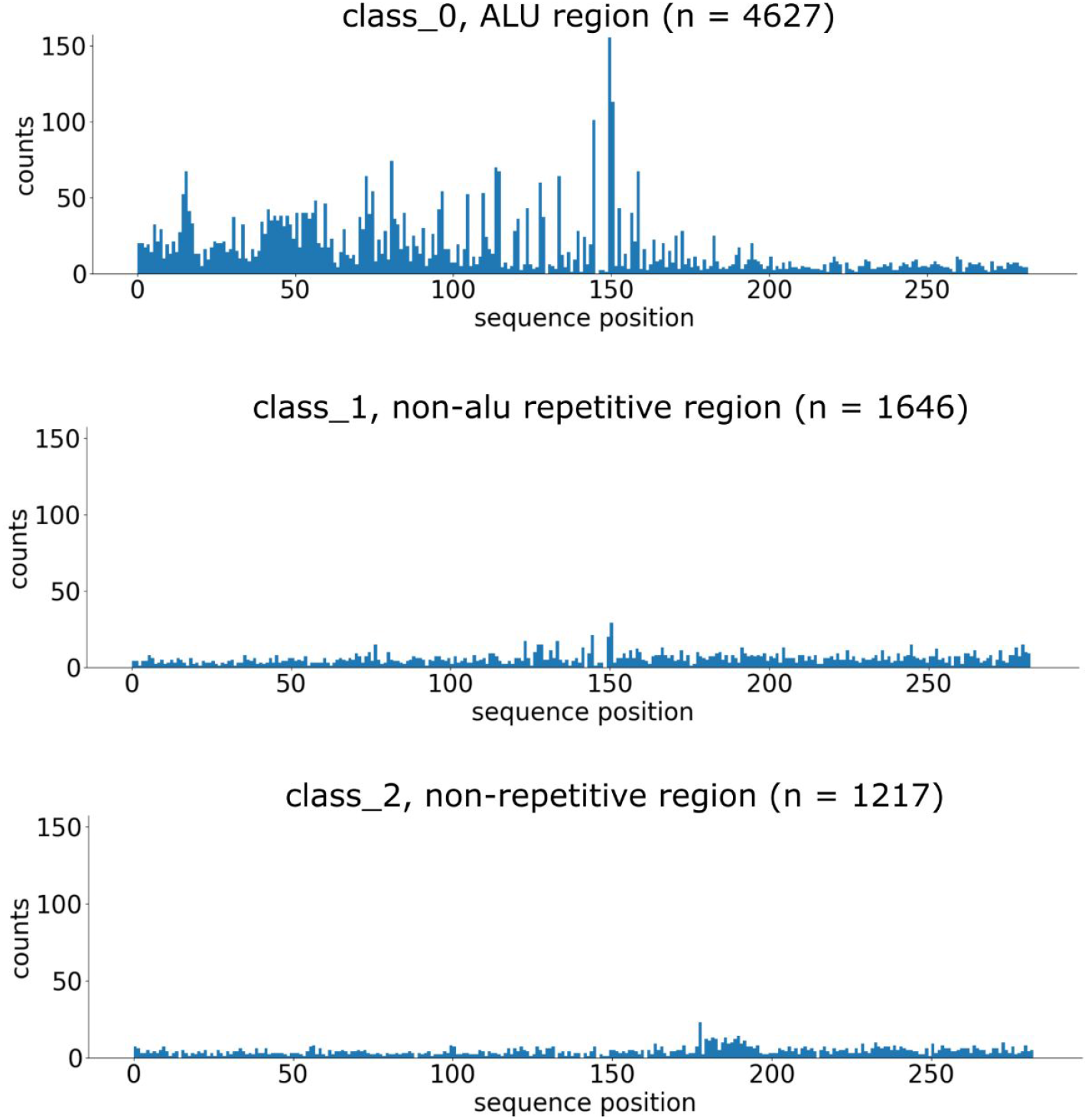
The histograms show the positional enrichment of a kernel (i.e. sequence positions at which subsequences leading to kernel activations higher than a threshold have been extracted from) for an RNA A-to-I editing classification data set consisting of three classes and input sequences of length 300. The motif corresponding to the kernel is shown in Figure 1C and is a known ALU repeat motif. The motif is therefore mainly found in the first class and preferentially starts close to sequence position 150. More details about the data set and the biological interpretation for this particular kernel can be found in the example workflow tutorial at github.

**Figure S2.**
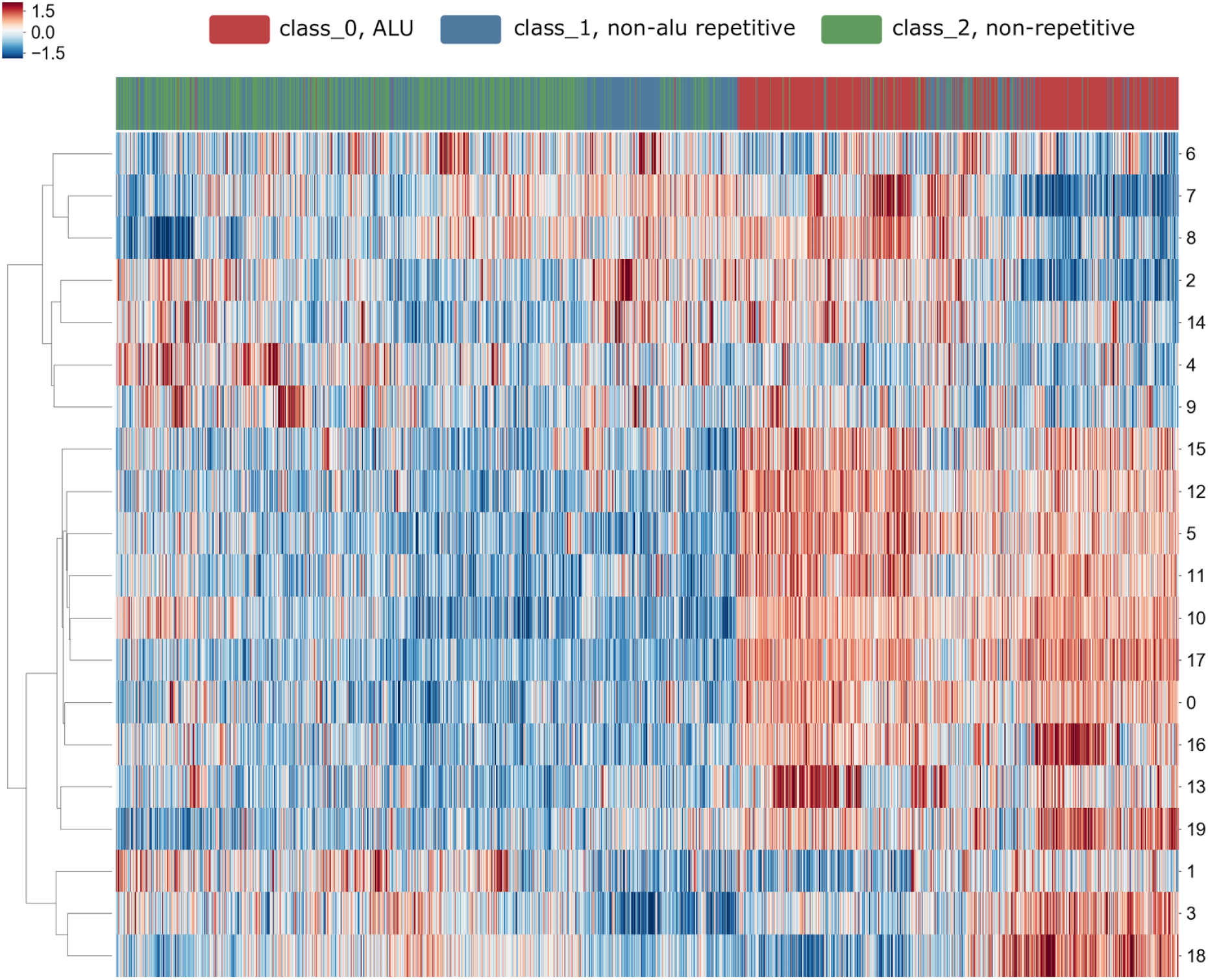
The heatmap shown here depicts a hierarchical clustering of both kernels (rows) and input sequences (columns) of a standardized matrix M where each cell M(i,j) represents the maximum activation value of kernel i for input sequence j. The resulting clustering is indicative of both class enrichment (i.e. kernels enriched or depleted in a certain class compared to the others), as well as motif co-occurrences. The heatmap shown here refers to the classification results on the RNA A-to-I editing data set showcased in the github tutorial. It is important to bear in mind that, besides putative motif co-occurrences, convolutional neural network kernels tend to learn strong motifs multiple times and hence, these tend to be clustered together. Nonetheless this visualization represents a valuable attempt to start looking for co-occurring motifs.

**Table S1.**
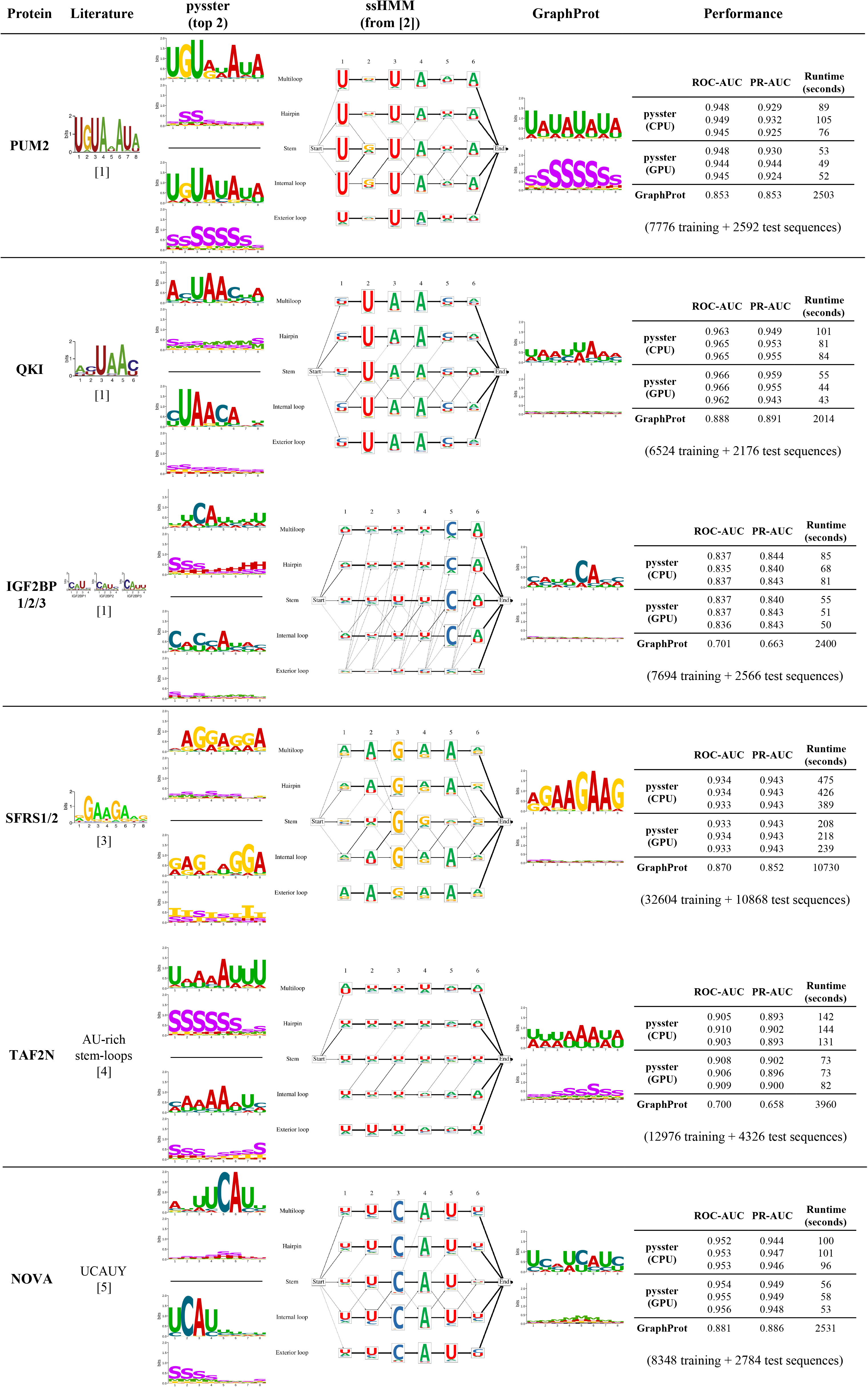
Performance comparison of pysster and GraphProt (v1.1.7) on CLIP-seq data for proteins with well-known RNA binding site motifs. For these binary classification tasks positive sets contained bindings sites from the protein of interest and negative sets contained randomly selected binding sites from 24 other proteins. All used sequences are of length 200 and centered at a binding site. Data are based on the ssHMM paper and the measurement source code and full output is available at https://github.com/budach/pysster/tree/master/tutorials/rbp. Aside from setting the motif length for GraphProt and the kernel length for pysster to 8 both tools were run with default parameters. Runtime includes RNA secondary structure prediction, model training and prediction of held-out test sequences and was measured on a machine with two E5-2697Av4 CPUs (32 physical cores) and an Nvidia Titan X GPU. Due to the non-deterministic nature of neural networks pysster was run three times to show that the predictive performance is stable. In addition, pysster performance is also shown for the CPU-only version. By default, pysster produces 30 motifs and the two with the highest importance score showing enrichment in the positive set are featured in the table. For ssHMM only the learned motifs are shown, as it is an unsupervised model only trained on the positive set.

